# eDNA-stimulated cell dispersion from *Caulobacter crescentus* biofilms upon oxygen limitation is dependent on a toxin-antitoxin system

**DOI:** 10.1101/2022.06.17.496608

**Authors:** Cecile Berne, Sébastien Zappa, Yves V. Brun

## Abstract

In their natural environment, most bacteria preferentially live as complex surface-attached multicellular colonies called biofilms. Biofilms begin with a few cells adhering to a surface, where they multiply to form a mature colony. When conditions deteriorate, cells can leave the biofilm. This dispersion is thought to be an important process that modifies the overall biofilm architecture and that promotes colonization of new environments. In *Caulobacter crescentus* biofilms, extracellular DNA (eDNA) is released upon cell death and prevents newborn cells from joining the established biofilm. Thus, eDNA promotes the dispersal of newborn cells and the subsequent colonization of new environments. These observations suggest that eDNA is a cue for sensing detrimental environmental conditions in the biofilm. Here we show that the toxin-antitoxin ParDE_4_ stimulates cell death in areas of a biofilm with decreased O_2_ availability. In conditions where O_2_ availability is low, eDNA concentration is correlated with cell death. Cell dispersal away from biofilms is decreased when *parDE_4_* is deleted, probably due to the lower local eDNA concentration. Expression of *parDE_4_* is positively regulated by O_2_ and the expression of this operon is decreased in biofilms where O_2_ availability is low. Thus, PCD by an O_2_-regulated toxin-antitoxin system stimulates dispersal away from areas of a biofilm with decreased O_2_ availability and favors colonization of a new, more hospitable environment.

## INTRODUCTION

Biofilms are multicellular communities attached to a surface, where complex exchanges and interactions occur between the different members. Biofilms first start with the attachment of individual bacterial cells to a surface and then grow into a more complex community when attached bacteria divide and new ones join. The biofilm lifestyle is considered beneficial for bacteria, as they usually provide shelters to xenobiotic stresses and predators and increase collective nutrient availability (Flemming *et al*., 2016). However, when conditions deteriorate, cells can leave the biofilm through a process referred to as dispersal, disseminate to new environments, and form new biofilms, enabling the colonization of new niches (Guilhen *et al*., 2017). Dispersal is triggered in response to various environment and biological cues, and understanding its regulation is important to determine how biofilms can be controlled.

Many Alphaproteobacteria use a strong polar adhesin to irreversibly attach to surfaces and form biofilms (Berne *et al*., 2015, Berne *et al*., 2018), with *Caulobacter crescentus* holdfast being the best characterized example. *C. crescentus* has a dimorphic life cycle, where each division cycle yields a sessile mother stalked cell and a motile daughter swarmer cell. Motile swarmer cells bear a single flagellum and multiple pili at the new pole. After the cell cycle progresses beyond a certain point, or upon contact with a surface, the newborn cells secrete a holdfast at the same pole and differentiate into stalked cells by retracting their pili, ejecting their flagellum, and synthesizing a thin cylindrical extension of the cell envelope called the stalk, which pushes the holdfast away from the cell body. While the chemical composition of the holdfast is not entirely elucidated, it is composed of polysaccharides with four different monosaccharide constituents, as well as DNA and peptide molecules of unknown nature (Merker & Smit, 1988, Hernando-Pérez *et al*., 2018, Hershey *et al*., 2019). Holdfast is an extremely strong bioadhesin (Tsang *et al*., 2006, Berne *et al*., 2013) crucial for irreversible cell adhesion to solid surfaces (Ong *et al*., 1990, Bodenmiller *et al*., 2004), colonization of air-liquid interfaces (Fiebig, 2019), and biofilm formation (Entcheva-Dimitrov & Spormann, 2004).

In some bacterial species, extracellular DNA (eDNA) plays a stabilizing role in the biofilm matrix (Okshevsky & Meyer, 2015, Campoccia *et al*., 2021). In contrast, we previously showed that *C. crescentus* eDNA produced via cell lysis negatively regulates biofilm formation and stimulates cell dispersal. eDNA binding to unattached holdfasts inhibits their adhesiveness, thereby inhibiting cell attachment to surfaces (Berne *et al*., 2010). In contrast, eDNA does not dislodge previously bound holdfasts. Therefore, eDNA prevents swarmer cells from joining mature biofilms but does not disperse existing biofilms. Because inhibition by eDNA is proportional to its concentration, we proposed that eDNA serves as a rheostat-like environmental cue to trigger dispersal when conditions are detrimental and cause cell death. However, it was not known if eDNA release is the simple consequence of random cell death occurring in the biofilm as conditions worsen or if it is the result of an active mechanism, such as programmed cell death (PCD) (Berne *et al*., 2010, Kirkpatrick & Viollier, 2010).

In this study, we demonstrate that cell death and eDNA release in a biofilm are regulated by a PCD mechanism that responds to oxygen availability. PCD in bacteria includes all genetically encoded mechanisms that lead to cell lysis (Lewis, 2000, Bayles, 2014). Toxin/Antitoxin systems (TAS) are important regulators of PCD (Rice & Bayles, 2008, Peeters & de Jonge, 2018). These systems are widespread in bacterial and archaeal genomes, but despite their abundance, the biological relevance of most TAS is still elusive (Fraikin *et al*., 2020). TAS are comprised of a stable toxin and its unstable antitoxin cognate. The antitoxin molecule usually antagonizes the toxin under “steady state” growth conditions; but, in PCD-triggering conditions, the antitoxin is inactivated, leading to an excess of free toxins that target key cellular processes in response to various environmental signals (Harms *et al*., 2018, Wang *et al*., 2020). There are currently eight types of TAS described in bacteria. The classification depends on the nature of the antitoxin (RNA in types I, III, and VIII, or small protein in the other TAS types), and the toxin (small protein in all but type VIII where the toxin is a small RNA), and how the antitoxin neutralizes the toxin activity (Song & Wood, 2020, Singh *et al*., 2021, Srivastava *et al*., 2021).

Among the 18 TAS identified in the *C. crescentus* genome (Ely, 2021), 13 have been studied experimentally, and belong to four different groups: 1) four paralogous RelBE (Fiebig *et al*., 2010) operons and one HigBA (Kirkpatrick *et al*., 2016) operon, belonging to the type II systems where the toxins (RelE or HigB) are known to be mRNA endonucleases; 2) four type II systems belonging to the ParDE family (Fiebig *et al*., 2010), where ParE toxins are usually DNA gyrase inhibitors; 3) three paralogs of HipBA, also a type II system, where the HipA toxins inhibit protein synthesis (Huang *et al*., 2020, Zhou *et al*., 2021); and 4) SocAB, the only member of the type VI TAS described so far, where the SocB toxin directly inhibits DNA replication (Aakre *et al*., 2013). The environmental conditions that trigger any of these TAS and their biological function are not yet fully identified.

In this study, we show that the ParDE_4_ TAS is involved in PCD and eDNA release in *C. crescentus* biofilms where it stimulates cell dispersal. We show that areas of a biofilm with decreased O_2_ availability experience more cell death. Cell viability is improved in a Δ*parDE_4_* mutant biofilm, especially in areas of decreased O_2_ availability, generating less cell lysis and less eDNA release. We also show that cell dispersal is decreased when *parDE_4_* is deleted, probably due to the lower local eDNA concentration. Expression of *parDE_4_* is positively regulated by O_2_ and the expression of this operon is decreased in biofilms where O_2_ availability is low. Thus, PCD by an O_2_-regulated TAS stimulates dispersal away from areas of a biofilm with decreased O_2_ availability.

## RESULTS

### The ParDE_4_ TAS is involved in biofilm inhibition and eDNA release of *C. crescentus* grown under static conditions

We previously showed that, in *C. crescentus,* eDNA is a cue that can trigger biofilm inhibition and dispersion by binding to holdfasts and reducing their adhesiveness. This mechanism is a result of cell lysis and eDNA release in the biofilm (Berne *et al*., 2010). To investigate if this eDNA release is the product of a specific PCD mechanism, we tested if a TAS was involved in promoting cell death in the biofilm, as previously suggested (Kirkpatrick & Viollier, 2010). If such a TAS is inactivated, one should observe less cell death, less eDNA release, and more biofilm formation. We examined the four ParDE-like and four RelBE-like individual in-frame deletion mutants previously described (Fiebig *et al*., 2010), as well as mutants lacking the four ParDE (“All *parDE ^-^* mutant”), the four RelBE (“All *relBE ^-^* mutant”) and the eight ParDE / RelBE operons (“All *parDE ^−^ All relBE ^-^*mutant”), for their ability to form biofilms compared to *C. crescentus* CB15 wild-type (WT). For these static biofilm assays, we grew cells in two ml plastic microfuge tubes sealed with AeraSeal breathable film, to allow for gas exchange, and incubated them statically at 30°C (Fig 1A). We defined these growth conditions as “moderate aeration”. All the tested mutants grew similarly to WT under these conditions (Fig. S1A).

**Figure 1:**
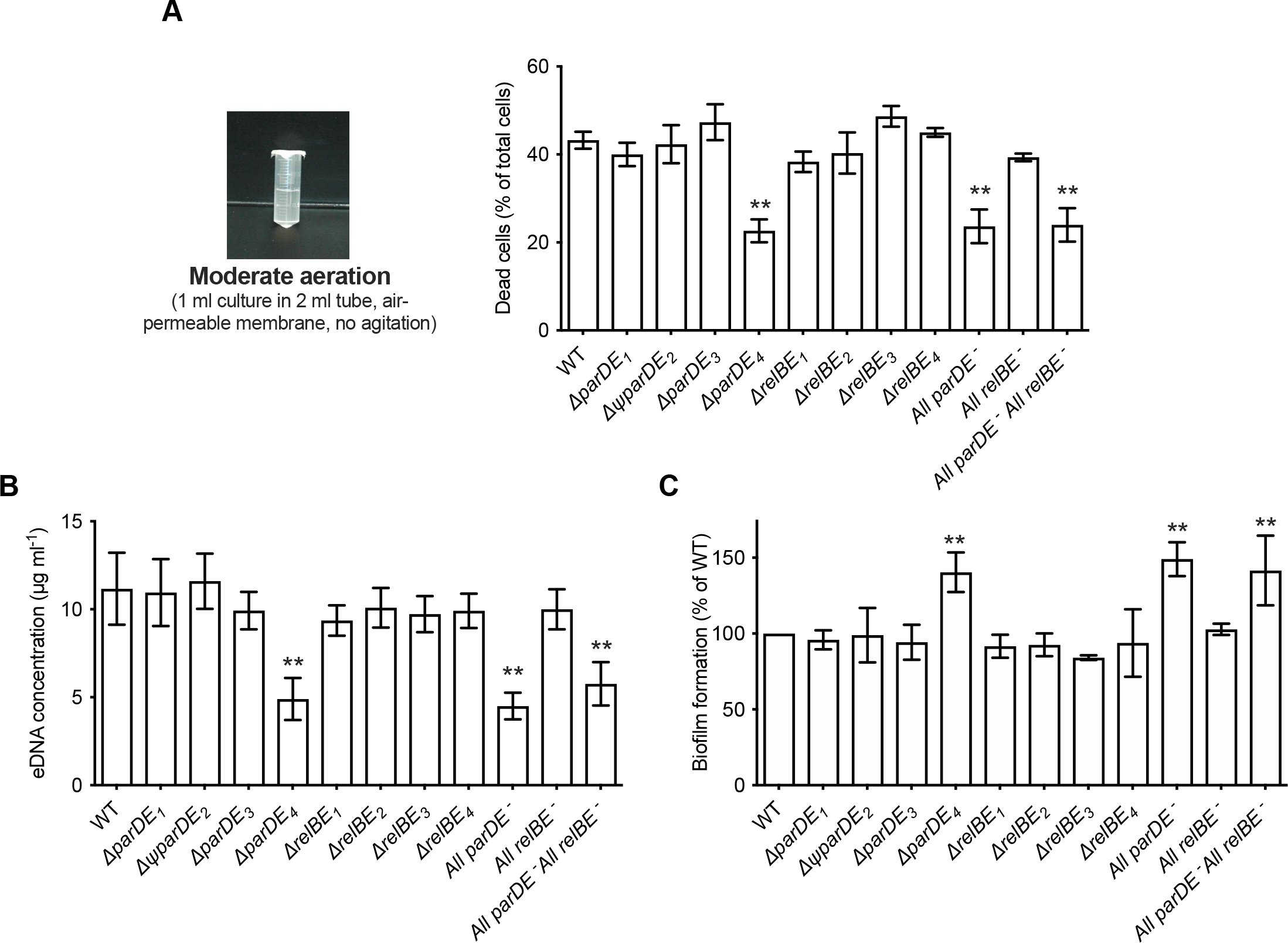
Role of the eight TAS in cell death, eDNA release, and biofilm formation. *C. crescentus* WT and the different TAS in-frame deletion mutants were grown for 48 h under moderate aeration conditions at 30°C, as depicted on the left. (A) Percentage of dead cells in the planktonic phase; results are expressed as a percentage of the total cells (live + dead) in the sample, quantified using the BacLight Live/Dead kit. (B) Quantification of eDNA released in the planktonic phase, quantified using PicoGreen reagent. (C) Biofilm formation, quantified by crystal violet staining; results are expressed as a percentage of biofilm formed compared to WT. Results are given as the average of four independent experiments, each run in duplicate, and the error bars represent the Standard Error of the Mean (SEM). Statistical comparisons are calculated using Student’s unpaired t-tests; only samples statistically different from WT are shown. ** *P* < 0.05.

We tested the ability of the TAS mutants to form biofilms under these growth conditions after 48 hours, and quantified cell death and eDNA release in these biofilms (Fig. 1). Among single mutants, Δ*parDE_4_* was the only strain that behaved differently compared to WT. It displayed increased viability in its biofilms and produced ∼30% more biofilm than the other strains (Fig. 1A and 1C). Furthermore, it released only about half of the amount of eDNA in the planktonic phase compared to WT (Fig 1B and S1B). The All *parDE ^−^* and the All *parDE ^−^* All *relBE ^-^* strains, where all four *parDE* operons and all *parDE* plus all *relBE* operons were deleted, respectively, behaved like the Δ*parDE_4_* single deletion mutant (Fig. 1). These results suggest that ParDE_4_ plays a role in biofilm regulation and eDNA release under our experimental conditions.

### The ParDE_4_ TAS plays a role in cell death in mature biofilms of *C. crescentus*

The *parDE_4_* operon is composed of the *parD_4_* antitoxin gene (CC2985 / CCNA_03080) and the *parE_4_* toxin gene (CC2984 / CCNA_03079), overlapping by 21 bp (Nierman *et al*., 2001, Fiebig *et al*., 2010, Marks *et al*., 2010). To assess the role of ParDE_4_ in cell death, eDNA release, and biofilm formation over time, we monitored biofilm formation on plastic strips grown under moderate aeration as depicted in Fig 1A. Over time, we also quantified eDNA release and cell death occurring in WT and Δ*parDE_4_* (Fig. 2). Cell death was reduced in Δ*parDE_4_* biofilms compared to WT, especially at longer time points when the biofilm reached maturation (Fig. 2 A and B). In addition, less eDNA was released in these mutant cultures (Fig. 2C). We also observed an increase in attached biomass in the Δ*parDE_4_* mutant (Fig 2D). These results support our previous findings that biofilm inhibition, eDNA release, and cell death are correlated (Berne *et al*., 2010). Furthermore, these results indicate that ParDE_4_ is involved in stimulating cell death and eDNA release in biofilms. Since eDNA stimulates dispersal from the biofilm (Berne *et al*., 2010), both the reduced cell death and eDNA release in the Δ*parDE_4_* mutant might contribute to the increased biofilm formation.

**Figure 2:**
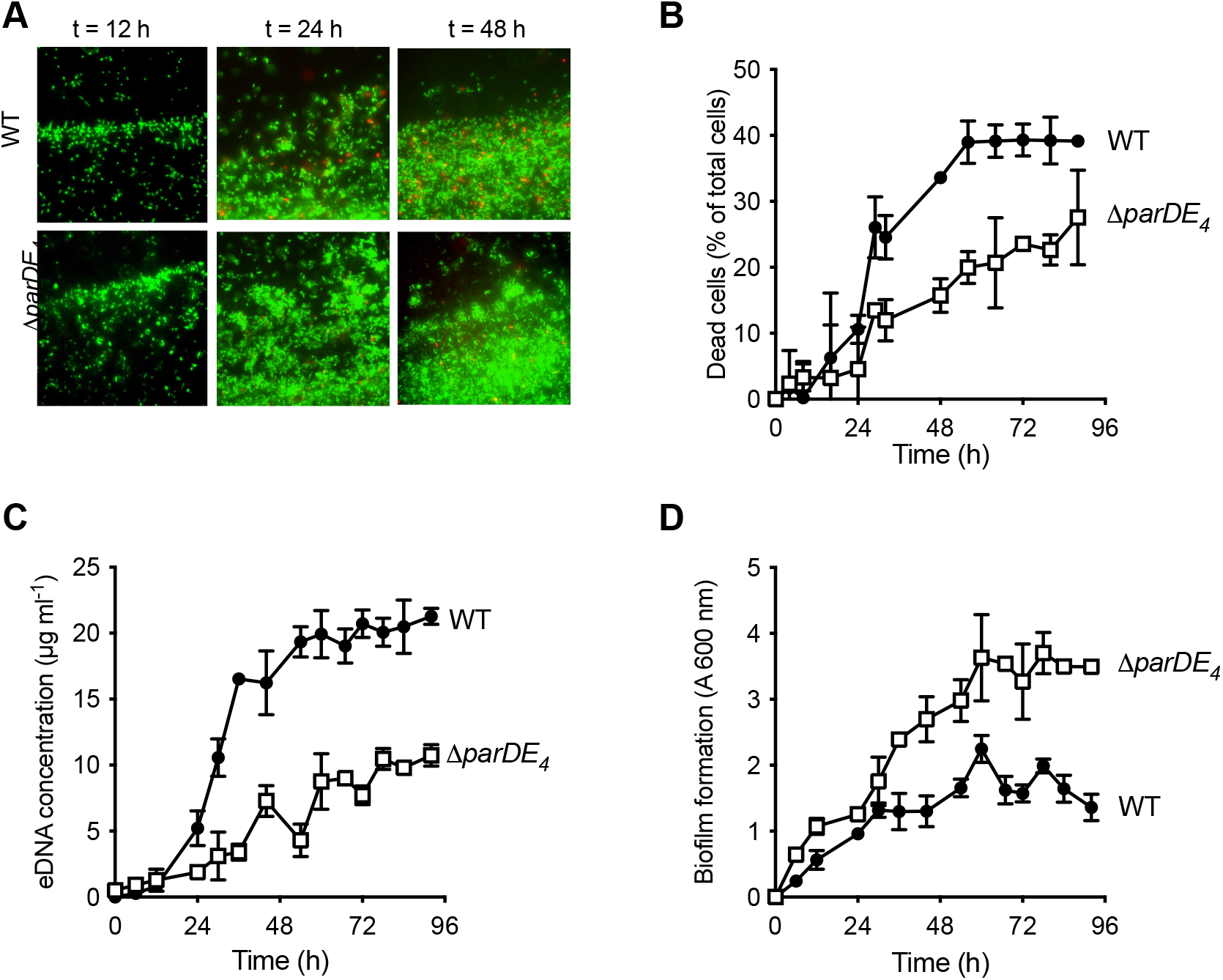
Involvement of the ParDE_4_ TAS in cell death, eDNA release, and biofilm regulation. (A) Biofilm formed on PVC strips stained with the BacLight Live/Dead reagent at different incubation times. Images represent overlays of the green (live cells) and red (dead cells) signals collected by epifluorescence microscopy. (B) Percentage of dead cells over time in the biofilm, calculated from BacLight Live/Dead stained cells using microscopy images. (C) eDNA release in the planktonic phase over time, quantified using PicoGreen staining. (D) Biofilm formation over time, quantified by staining the attached biomass with crystal violet. *C. crescentus* WT and Δ*parDE_4_* are represented by solid circles and open squares symbols respectively. The results are given as the average of three independent experiments, each run in duplicate, and the error bars represent the SEM.

### The ParD_4_ antitoxin protects against cell death in the biofilm

In TAS, cell death usually occurs when there is an imbalance between the amount of stable toxins and labile antitoxins produced in the cell (Harms *et al*., 2018). To assess the role of ParD_4_ antitoxin expression, we expressed it using the low copy vanillate-inducible replicating plasmid pMT630 (Thanbichler *et al*., 2007) and monitored biofilm formation and eDNA release when the antitoxin is constitutively expressed. When we constitutively expressed the ParD_4_ antitoxin in WT, biofilm formation was increased and eDNA concentration was decreased (Fig. 3A). However, in a Δ*parE_4_* mutant lacking the ParE_4_ toxin, there was no effect of *parD_4_* expression on biofilm formation and eDNA concentration (Fig. 3B), showing that 1) ParD_4_ has a protective effect against cell death in the biofilm, and 2) this effect depends on the presence of the toxin ParE_4_.

**Figure 3:**
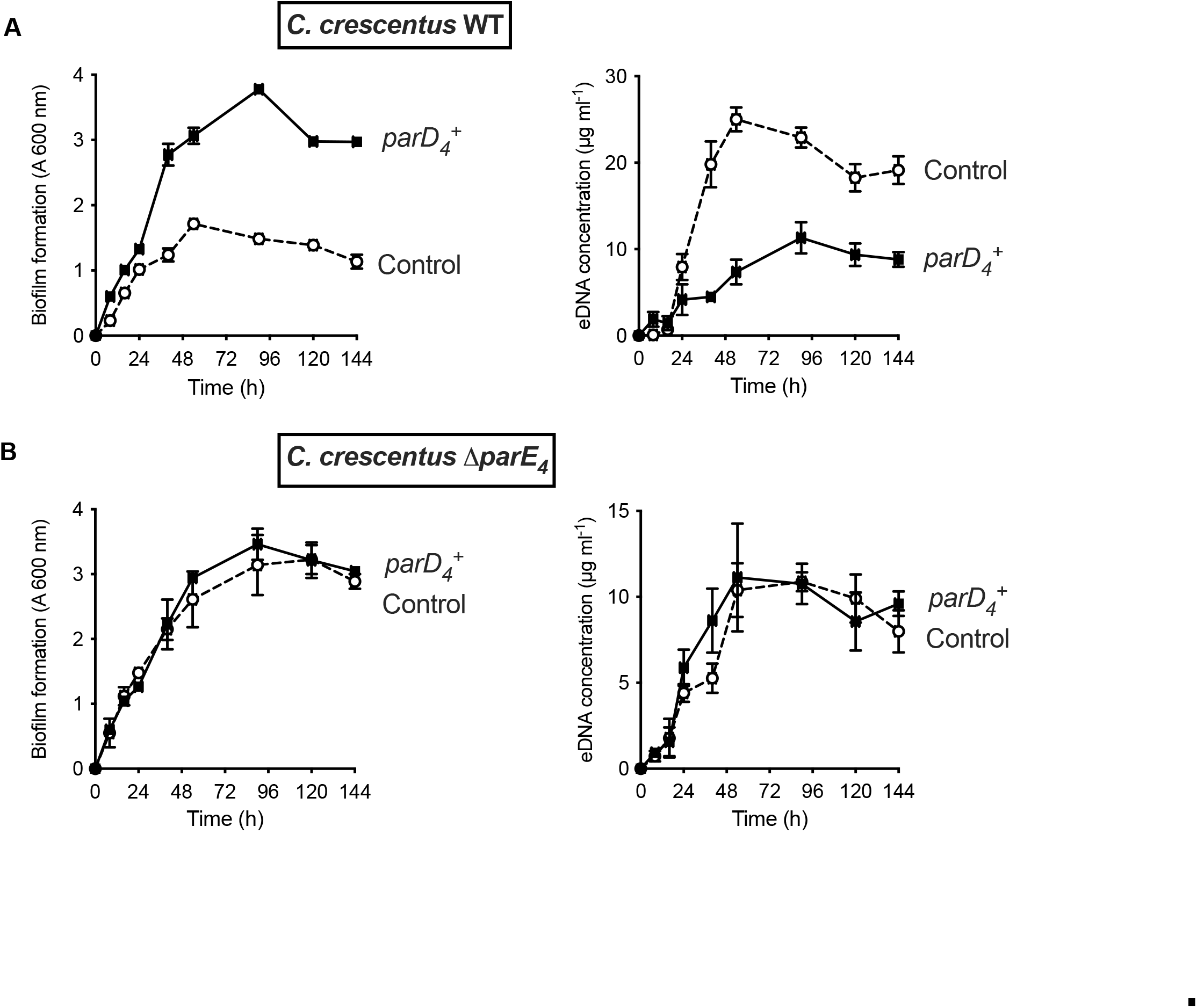
Effect of *parD_4_* antitoxin gene induced constitutive expression on biofilm formation and eDNA release. The *parD_4_* was cloned into the low copy replicating pMT630 plasmid and expressed using the P*van* promoter (induction using 500 µM vanillate). Biofilm formation (left panels) and eDNA release (right panels) for strains expressing the *parD_4_* antitoxin gene (solid square symbols) or bearing the empty pMT630 plasmid (open circle symbols) in WT (A) and Δ*parE_4_* (B).

Next, we wanted to determine if the behavior of the Δ*parDE_4_* mutant was due to lack of cell death and eDNA release, or also possibly to an impaired response to eDNA biofilm inhibition in the Δ*parDE_4_* mutant. We tested how WT and Δ*parDE_4_* behaved in the presence of exogenous eDNA by monitoring the amount of biofilm formed in the presence of spent media from cultures of different strains. We showed previously that eDNA present in spent medium from saturated cultures inhibits biofilm formation (Berne *et al*., 2010). Spent media, containing various concentrations of eDNA from cultures of either WT or Δ*parDE_4_* grown to late stationary phase were used in biofilm assays, as done previously (Berne *et al*., 2010). Because Δ*parDE_4_* produces less eDNAthan WT (∼8 µg/ml compared to ∼12 µg/ml of eDNA present in the spent medium of a saturated culture for Δ*parDE_4_* and WT respectively, see Fig. S2), we first determined the amount of eDNA present and added an appropriate amount of spend medium to obtain the same final concentration of eDNA. The amount of biofilm formed by WT and Δ*parDE_4_* were similar for the same total amount of eDNA (Fig. S2), showing that, when exposed to the same amount and source of eDNA, both WT and Δ*parDE_4_* form similar amounts of biofilms and that eDNA from WT or Δ*parDE_4_* have the same biofilm inhibitory activity. Furthermore, the inhibition response is positively correlated with the amount of eDNA present in the spent medium in a similar manner for both strains (Fig. S2), in agreement with our previous results (Berne *et al*., 2010). Therefore, the increase in biofilm formation by the Δ*parDE_4_* mutant is not due to an impaired response to eDNA but is likely due to less cell death causing a more rapid mass increase and/or to less cell dispersal stimulation by eDNA.

### ParDE_4_ promotes population dispersal in the biofilm

To understand the dynamics of biofilm formation in the WT and Δ*parDE_4_* strains, we grew a mixed culture of differently fluorescently labeled WT and Δ*parDE_4_* at a 1:1 ratio in flow cells. The evolution of each population in the biofilm was monitored over time (Fig. 4A). Patterns in the spatial organization of the biofilm could be observed at early stages of biofilm maturation (Fig. 4A), with formation of homogenous microcolonies due to clonal growth. This observation is in agreement with previous reports of *C. crescentus* biofilm growth in flow cells (Entcheva-Dimitrov & Spormann, 2004, Rossy *et al*., 2019). While in the early stages of biofilm formation the ratio of WT and Δ*parDE_4_* was maintained, the mutant population rapidly outcompeted the WT at later stages. After 96 h, around 80% of the attached bacteria were Δ*parDE_4_* (Fig. 4B).

**Figure 4:**
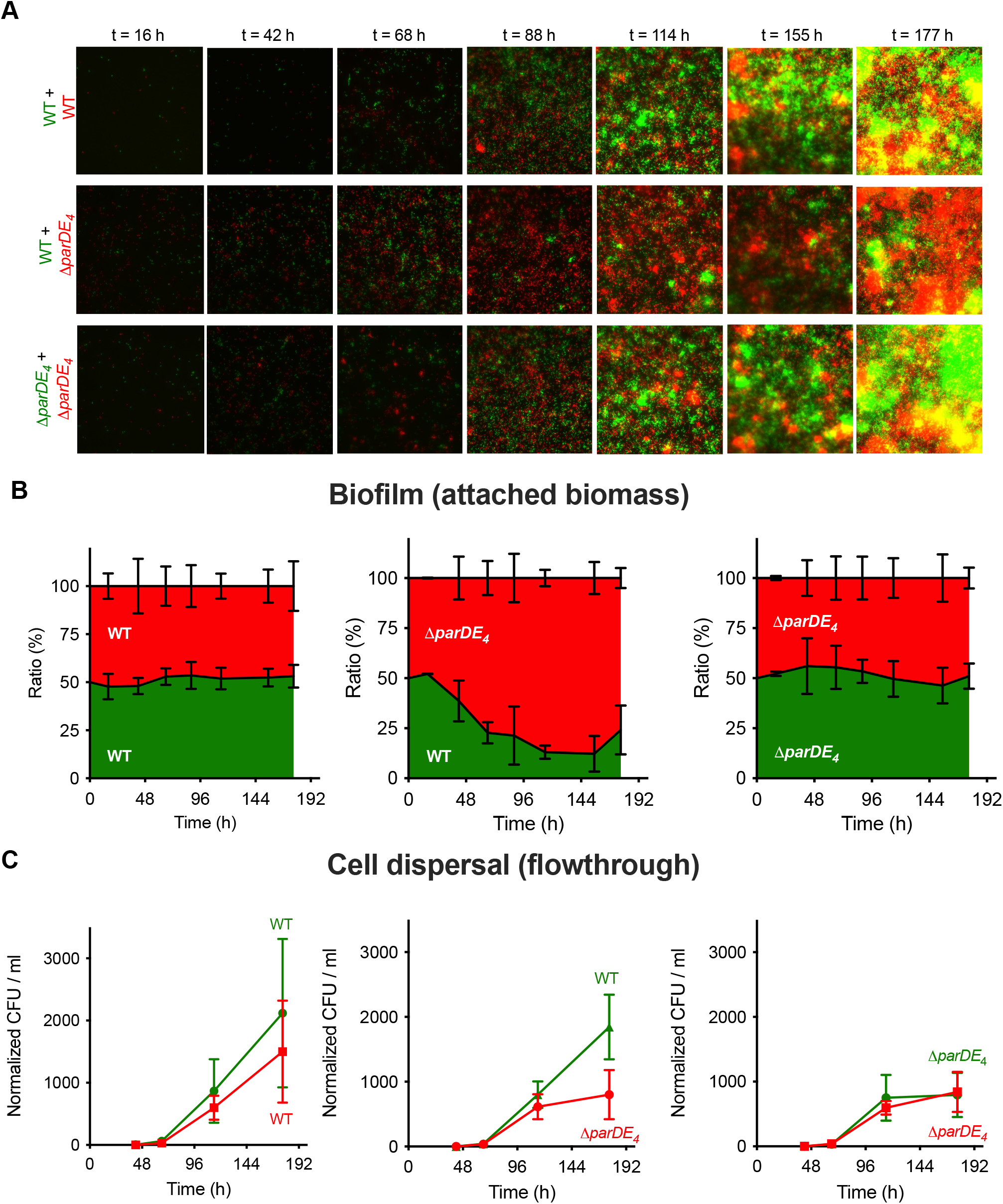
Biofilm formation and dispersion in mixed cultures. Differentially fluorescently tagged populations of WT and Δ*parDE_4_* were mixed to a 1:1 ratio and grown in flow-cells over time. (A) Representative fluorescence microscopy images of mixed culture biofilms grown in flow cells. One population is represented in green and the other one in red. (B) Ratio of each population over time. Results are given as a percentage of total fluorescent area (green + red) representing both populations, calculated from microscopy images (average of 10 fields of view per time point, run in triplicate during two independent experiments where fluorescent markers were swapped). (C) Cell dispersal measured as Colony Forming Units (CFU ml^-1^) released in the flowthrough downstream of the flow-cells. Results are normalized to the number of colonies measured for WT at t = 42 h (beginning of the experiment). Results are expressed as an average of 2-4 serial dilution counts of 3 samples per time point. Flow-cells were run in triplicate for each of the two independent experiments where fluorescent markers were swapped. Errors bars represent the SEM.

To test the dispersal rate of both strains, we measured dispersal from the biofilm by quantifying the number of cells released from the biofilm over time in the flow-cells. This was done by collecting flow-through samples downstream of the flow cell and quantifying the number of single cells released from each population (Fig. 4C). Surprisingly, while there was more biomass of Δ*parDE_4_* cells than WT in the mixed biofilm experiment, there were more WT cells released over time compared to Δ*parDE_4_*, indicating that the dispersal of WT is more efficient (Fig. 4C).

In summary, these results combined with those of previous sections suggest that the observed increased biofilm formation in the Δ*parDE_4_* is due to a combination of increased attached biomass because of reduced cell death and decreased dispersion efficiency.

### The ParDE4 response is correlated with O_2_ availability

Previous work investigated the regulation of the *parDE_4_* operon as a function of O_2_ availability on planktonic, exponentially growing cells (Fiebig *et al*., 2010). Transcriptome profiling by microarray performed under hypoxic stress (limiting O_2_ conditions) showed a two-fold decrease in *parD_4_* expression, albeit not statistically significant (Fiebig *et al*., 2010). In our hands, PCD triggered by the ParDE_4_ TAS is more pronounced when the biofilm reaches maturation (Fig. 2). A number of environmental changes occur as the biofilm matures, including variation of O_2_ availability. Since *C. crescentus* is an obligate aerobe, O_2_ depletion could be a major detrimental factor, triggering cell death and dispersal, as is the case in other species (Thormann *et al*., 2005, McDougald *et al*., 2012).

In order to test the effect of O_2_ on ParDE_4_-mediated cell death, we grew cells with vigorous shaking (“maximal aeration”) as compared to the non-shaking “moderate aeration” condition (Fig. 5A). Interestingly, a decrease in Δ*parDE_4_* mediated eDNA release did not occur in cells grown under vigorous shaking, (“maximal aeration” conditions) compared to the “moderate aeration” conditions (Fig. 5B and S1), suggesting that ParDE_4_ is not active under those conditions. To determine if ParDE_4_ expression is regulated by aeration conditions, we monitored its transcription using a *lacZ* fusion under maximal aeration growth compared to growth under moderate aeration conditions. To validate those growth conditions as providing different O_2_ availability, we used a *lacZ* fusion to the promoter of *ccoN*, encoding the cytochrome *cbb*_3_ oxidase subunit I. This gene is highly expressed when *C. crescentus* cells experience limiting O_2_ levels (Crosson *et al*., 2005) and its expression can be used as a biosensor to monitor O_2_ availability. P*_ccoN_* expression was 10-13 times more active under moderate aeration conditions (Fig. 5C), confirming that O_2_ availability is limited under those growth conditions. We found that P*_parDE4_* transcription was approximately 2-3 times higher under maximal aeration growth (Fig. 5C), suggesting that ParDE_4_ expression is regulated by O_2_ availability.

**Figure 5:**
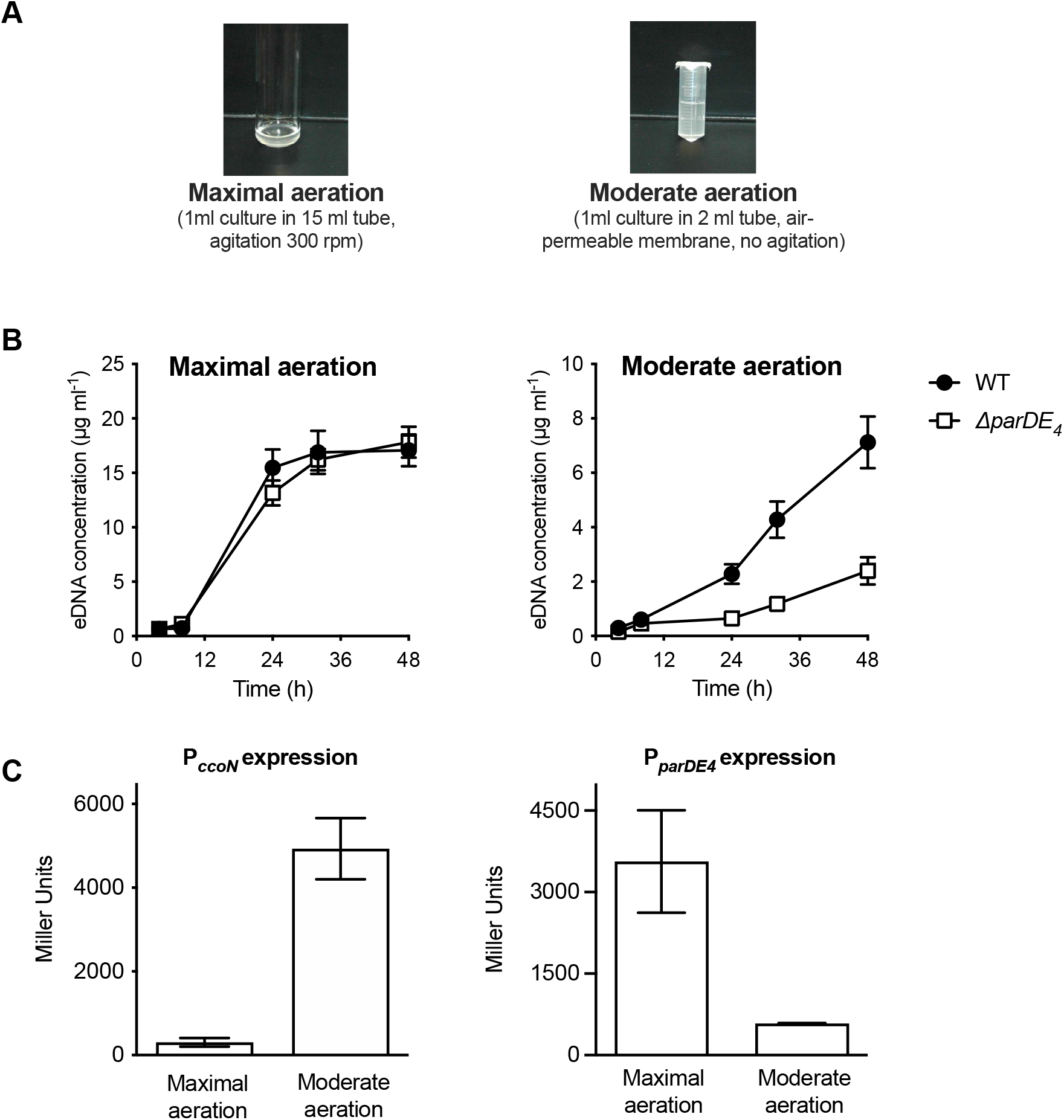
P*parDE_4_* expression is induced under maximal aeration growth conditions. (A) Images of cultures grown under conditions providing different amounts of O_2_, termed maximal and moderate aeration, respectively. (B) Quantification of eDNA released in the planktonic phase of WT (solid circles) and Δ*parDE_4_* (open squares), quantified using PicoGreen staining. Results are given as the average of five independent experiments and the error bars represent the Standard Error of the Mean (SEM). (C) ß-galactosidase activity of P*_parDE4_*-*lacZ* (right) and P*_ccoN_*-*lacZ* transcriptional fusions in WT grown under maximal and moderate aeration conditions (as illustrated in panel A). The results represent the average of 6 independent cultures (assayed on 3 different days) and the error bars represent the SEM.

Since the above results suggested that O_2_ availability is important for ParDE_4_- controlled biofilm regulation, we tested two additional growth conditions to obtain an intermediate level of O_2_ (“high aeration”) and an hypoxic level (“limited aeration”) (Fig. 6A), as assessed by measuring the expression of P*_ccoN_* (Fig. 6B). We first confirmed that the expression of P*_parDE4_* is inversely correlated to the O_2_ level in the cultures (Fig. 6B). We then compared the ratio of eDNA released in Δ*parDE_4_* relative to WT in the different aeration conditions. We found that there was twice as much eDNA released by WT compared to Δ*parDE_4_* under conditions where O_2_ levels are the most reduced whereas the levels were similar under maximal aeration (Fig. 6C). Concomitant with the decrease in eDNA release, we saw that biofilm formation by the Δ*parDE_4_* increased relative to WT as O_2_ availability decreased (Fig. 6D). In summary, lower O_2_ availability correlated with less eDNA release and more biofilm formation by the Δ*parDE_4_* strain compared to WT. We therefore conclude that ParDE_4_ triggers cell death when O_2_ is limited, which in turn releases eDNA. As shown in the previous sections and in our previous work (Berne *et al*., 2010), eDNA inhibits biofilm formation.

**Figure 6:**
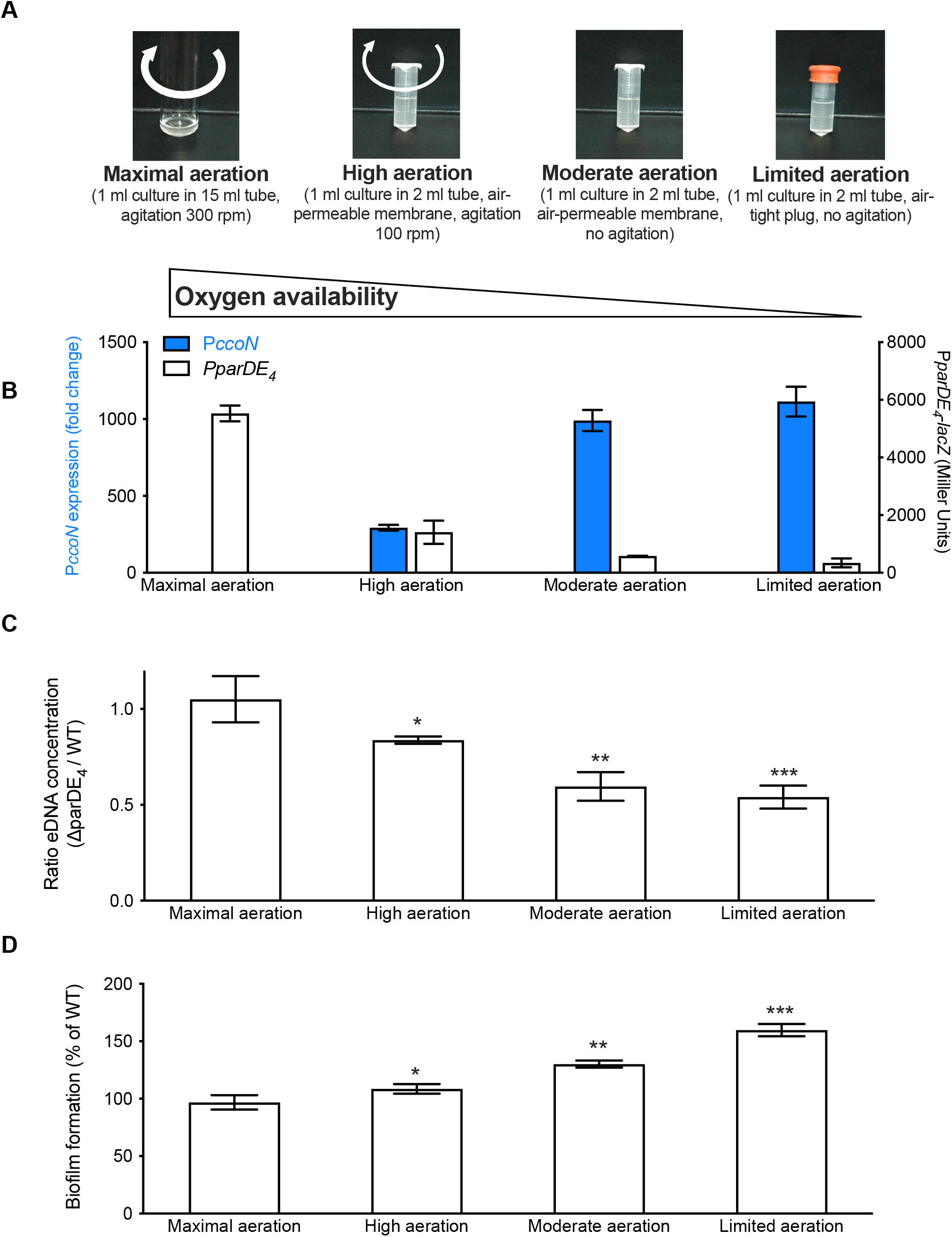
eDNA release and biofilm formation under variable O_2_ availability. (A) Images of cultures providing different amounts of O_2_, termed maximal, high, moderate, and limited aeration, respectively. (B) Assessment of P*_parDE4_* and P*_ccoN_* expression (white and blue bars respectively) by measuring ß-galactosidase activity of the P*_parDE4_*-*lacZ* transcriptional fusions in WT, and transcription of *ccoN* relative to *rpoD* as a function of various aeration conditions by RT-qPCR. (C) Ratio of the eDNA release concentration measured in Δ*parDE*_4_ planktonic phase over WT, in cultures grown under the different aeration conditions. Results are given as the calculated ratio of the eDNA concentration measured in 6 independent replicates, each run in duplicate, and error bars represent the SEM. Statistical comparisons to “maximal aeration” conditions are calculated using Student’s unpaired t-tests;* *P* < 0.5, ** *P* < 0.05, *** *P* < 0.005. (D) Biofilm formation after 24 hours when cells are grown under the different aeration conditions. Results are given as the average of 3-5 independent replicates, each run in triplicate, and error bars represent the SEM. Statistical comparisons to “maximal aeration” conditions are calculated using Student’s unpaired t-tests;* *P* < 0.5, ** *P* < 0.05, *** *P* < 0.005.

Next, we wanted to determine if O_2_ limitation and *parDE_4_* expression are correlated at the single cell level within a population. For that purpose, we created a fluorescent reporter fusion for the *parDE_4_* operon by fusing its promoter region to a promoterless GFP construct to monitor *parDE_4_* expression via GFP fluorescence signal. We also constructed a mCherry fusion to the promoter region of *ccoN* to monitor O_2_ availability for each cell using mCherry fluorescence as a proxy. We grew cells carrying both reporters under our four growth conditions with different O_2_ availability (Fig. 6A). We first quantified the number of GFP and mCherry expressing cells under each condition (Fig. 7A). As expected from the activity of the *ccoN* promoter, the number of mCherry expressing cells decreased as growth conditions became less anoxic: while more than 60% of cells expressed mCherry under limited aeration, only 6% did so under maximal aeration (Fig. 7A). In contrast, we observed an opposite trend for GFP expression driven by the *parDE_4_* promoter; it increased with increasing O_2_ availability in the culture, with more than 40% of cells expressing GFP under maximal and high aeration conditions and around 10% under moderate and limited aeration (Fig. 7A). We also monitored the red and green fluorescence intensity per cell in the four different growth conditions (Fig 7B and S3). There was a strong anticorrelation between the expression of *ccoN* and *parDE_4_* in single cells: cells with high expression of one of the two promoters rarely expressed the other. In other words, cells which experience hypoxia rarely express *parDE4*, while cells sensing higher levels of O_2_ levels do, and reciprocally. These results confirm what we previously observe at the population level, *i.e.* that *parDE_4_* expression is correlated with O_2_ availability.

**Figure 7:**
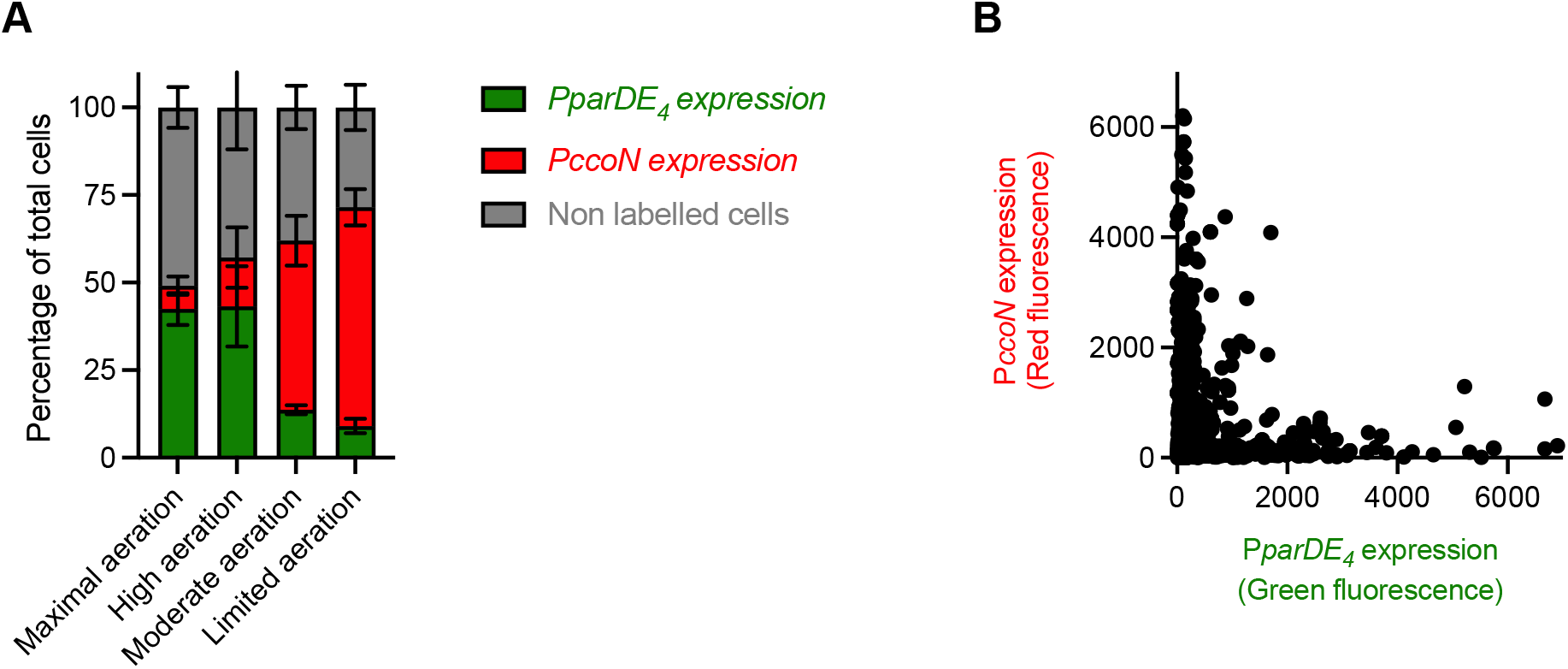
Expression of the *parDE_4_* operon is anticorrelated with a hypoxia reporter. (A) Number of cells expressing GFP (P*parDE_4_* expression) and mCherry (P*ccoN* expression) in the whole population when cells were grown under maximal, high, moderate, and limited aeration, providing different amounts of O_2_. (B) Red and green fluorescence intensity of single cells grown under maximal, high, moderate, and limited aeration (all conditions combined). WT cells carrying both pMR20-P*parDE_4_-gfp* and pMR10-P*ccoN-mcherry* plasmids were grown to OD_600_ = 0.4-0.6 under maximal, high, moderate, and limited aeration (Fig. 6A) and imaged by epifluorescence microscopy. More than 5,000 cells from at least three independent replicates were quantified for the number of cells with a green or red fluorescent signal (A) and the intensity of these fluorescent signals (B).

### ParDE_4_ is required for increased cell death in biofilm areas with lower O_2_ availability

To assess how local amounts of O_2_ might influence ParDE_4_ controlled cell death within an established biofilm, we next assessed the pattern of cell death and *parDE_4_* expression in different areas of the biofilm. We placed a sterile microscopy-grade clear polyvinyl chloride (PVC) strip inside Aeraseal covered microtubes. We monitored 3 distinct areas: 1) the air-liquid interface area at the meniscus (highest air exchange); 2) the middle of the strip (moderate air exchange); and 3) the bottom of the strip (limited air exchange) (Fig. 8). We determined the *parDE_4_* promoter activity using a P*_parDE4_*-*lacZ* transcriptional fusion and fluorescein Di-β-D-Galactopyranoside (FDG), a fluorogenic substrate whose fluorescence is turned on by cleavage by β-galactosidase (Rotman *et al*., 1963). The fluorescent signal decreased in cells attached at the bottom of the biofilm compared to the meniscus or the middle of the strip, indicating that the P*_parDE4_*-*lacZ* reporter is less active in areas with lower O_2_ availability (Fig. 8A).

**Figure 8:**
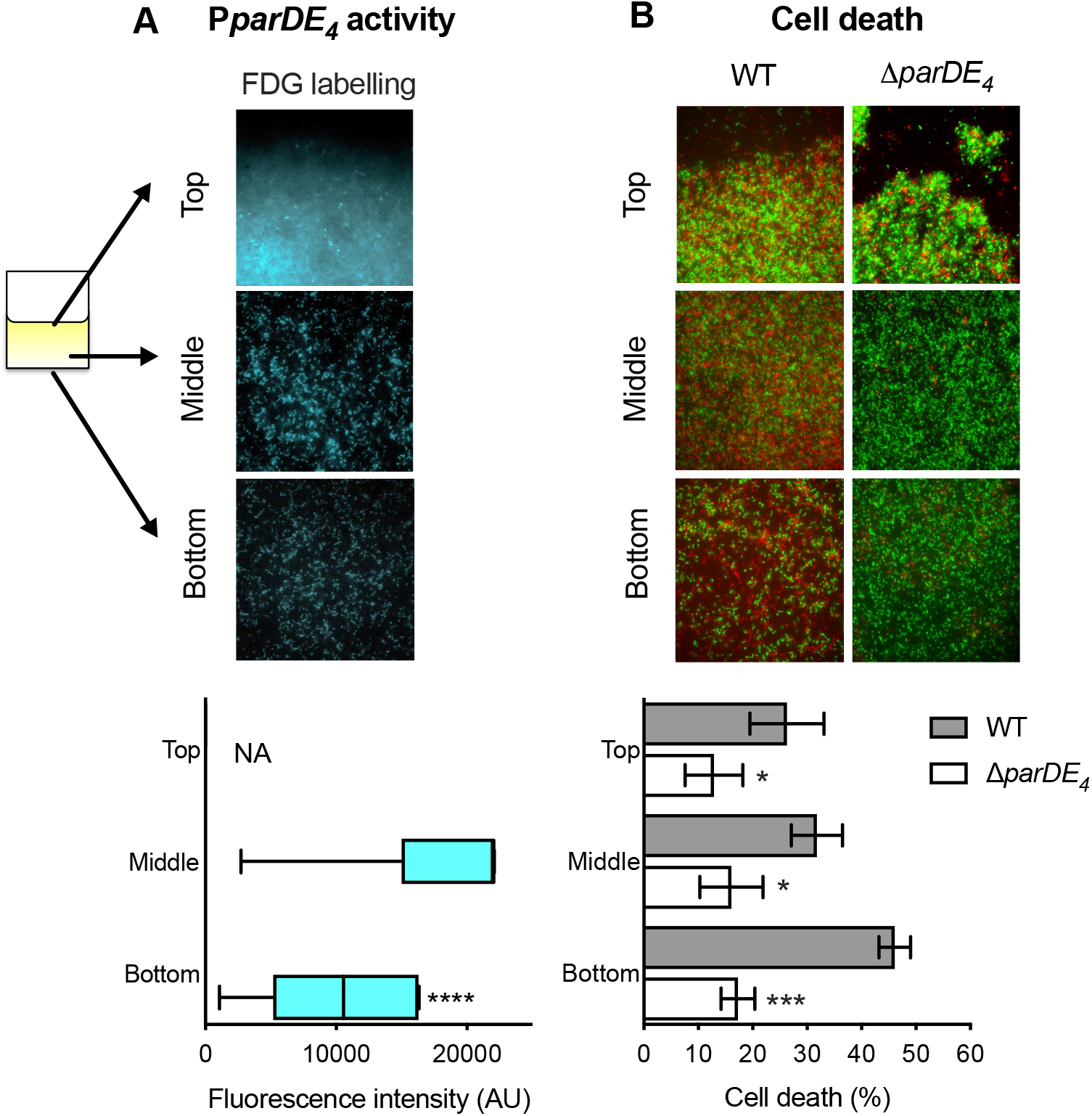
Spatial regulation of *parDE_4_* expression and ParDE_4_-mediated cell death in the biofilm. Biofilms were grown on PVC strips. Three locations were monitored: the air-liquid interface (Top); the middle of the strip (Middle) and the bottom of the strip (Bottom). (A) P*parDE_4_* activity in cells attached at the three different locations. WT cells carrying the P*parDE_4_-lacZ* plasmid were grown for 36 h, and *PparDE_4_* activity was measured by the amount of fluorescein cleaved by the ß-galactosidase from the fluorogenic FDG substrate. Fluorescence intensity was quantified using microscopy images of 10 fields of views of three independent replicates and results are shown as min to max box and whisker plots. Statistical comparisons are calculated using Mann-Whitney unpaired t-tests. **** *P* < 0.0001. We could not determine the amount of FDG hydrolyzation at the air-liquid interface, as the biofilm was too dense to accurately quantify the fluorescent signal, due to saturation. (B) Cell death quantification of biofilms of WT and Δ*parDE_4_* after 36 h. Percentage of dead cells over time in the biofilm, calculated from BacLight Live/Dead stained cells using microscopy images. Results are given as the average of percentage of dead cells (red) in 10 fields of views of three independent replicates and are shown in gray and white bars for WT and Δ*parDE_4_* respectively. Error bars represent SEM. Statistical comparisons to WT in the same condition are calculated using Student’s unpaired t-tests;* *P* < 0.5, *** *P* < 0.005.

When we monitored cell death in WT and Δ*parDE_4_* biofilms grown in the same manner, we found that WT displayed more cell death away from the meniscus, showing that cell death preferentially occurs where O_2_ is limited (Fig. 8B). Indeed, twice as many cells were dead at the bottom of the strip compared to the top (Fig. 8B). However, in the Δ*parDE_4_* mutant, the amount of cell death was not significantly different between the distinct locations within the biofilm, indicating that the ParDE_4_ TAS regulates programmed cell death induced by O_2_ limitation.

## DISCUSSION

While eDNA has been shown in the past to be a major component of the biofilm matrix that plays a role in overall biofilm architecture for different species (Okshevsky & Meyer, 2015, Campoccia *et al*., 2021), we previously reported a novel role for this molecule, as an environmental cue to sense deleterious environments in a biofilm and promote cell dispersion in *C. crescentus* (Berne *et al*., 2010). We showed that cell death and eDNA release increase during biofilm formation and that eDNA prevents newborn swarmer cell from attaching to surfaces and settling into a biofilm by binding specifically to and inhibiting the adhesiveness of the holdfast. Because stalked cells attached by their holdfast cannot be dislodged from a surface by eDNA, this mechanism promotes swarmer cell dispersal without causing a potentially undesirable dissolution of the existing biofilm. While our previous results support the role of eDNA as a cue that prevents settling of swarmer cells (Berne *et al*., 2010, Kirkpatrick & Viollier, 2010), it was not known if this mechanism was simply a consequence of random cell death or the result of an active mechanism.

In this work, we show that *C. crescentus* biofilms experience increasing cell death in regions of a mature biofilm where O_2_ becomes limiting. We show that this increased cell death in biofilm regions of O_2_ limitation depends on the *parDE_4_* TAS. We also show that *parDE_4_*-dependent cell death is highest in conditions where O_2_ is limited. Under conditions of maximal aeration, cell death and eDNA release are equivalent in WT and in the *parDE_4_* mutant, whereas they are higher in WT under conditions of limited aeration.

Interestingly, even if cells of the *parDE_4_* mutant increase their biofilm biomass more rapidly than WT, they have a reduced dispersal from the biofilm. This result is consistent with our previous findings that eDNA release increases as biofilms mature and that eDNA inhibits swarmer cell adhesion by binding to their holdfast, but does not dislodge previously bound cells (Berne *et al*., 2010). We suggest that, as the *C. crescentus* biofilm increases in biomass, O_2_ limitation activates the ParDE_4_ TAS, resulting in cell death and concomitant eDNA release, which promotes dispersal of swarmer cells by binding to their holdfast and inhibiting their adhesion. This process enables the colonization of new environments while preserving the survival of the initial biofilm (Fig. 9).

**Figure 9:**
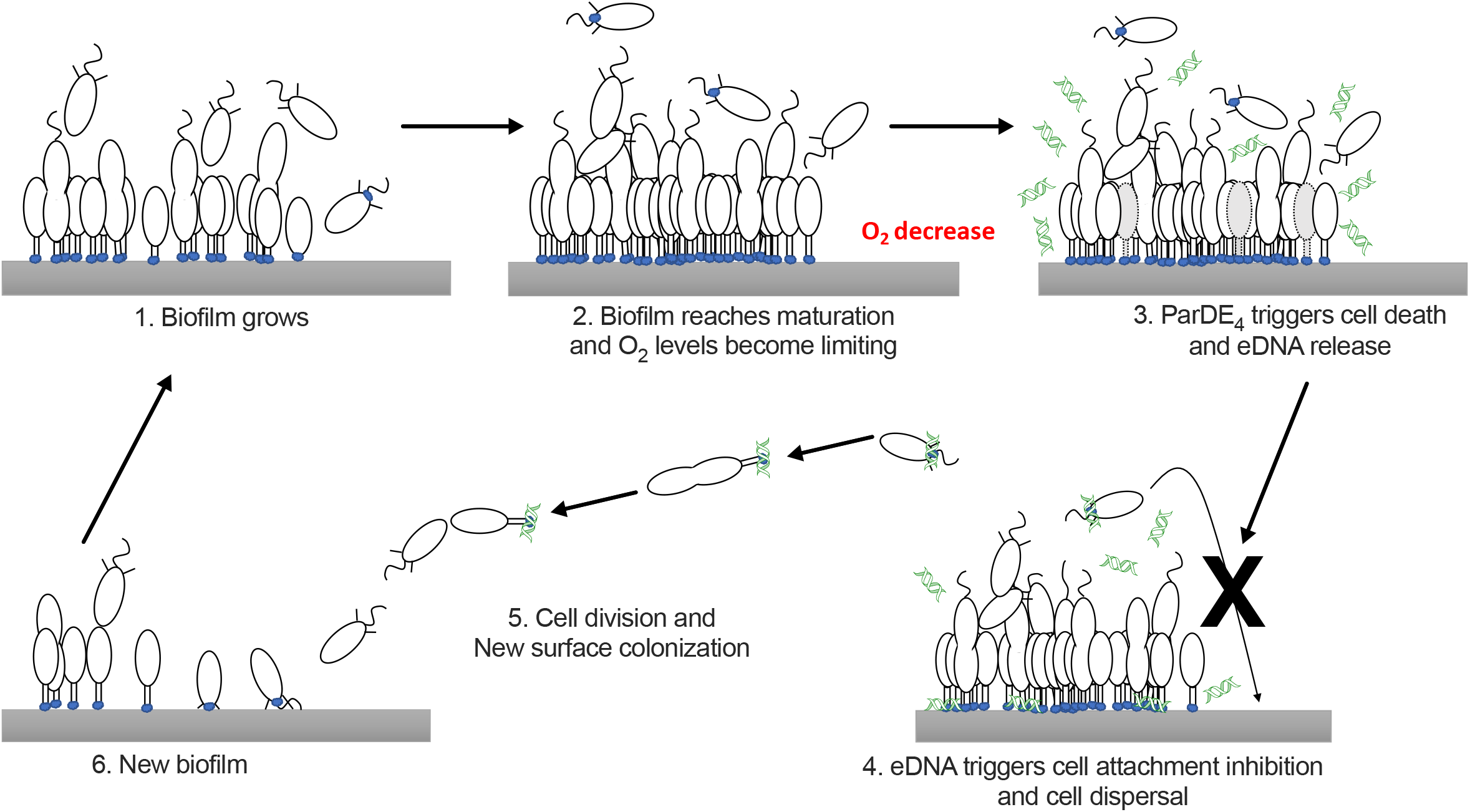
Schematic representation of the hypothetical mechanism of ParDE_4_-dependent regulation of cell death and dispersal upon O_2_ limitation. Once the biofilm reaches maturation, O_2_ availability becomes limiting. ParDE_4_-mediated PCD is initiated upon O_2_ deprivation and targeted cells release eDNA via cell lysis. eDNA specifically binds to holdfasts, preventing new cells from attaching, but does not influence already attached cells. Unable to join the biofilm, swarmer cells disperse, divide, and their offspring eventually find a new surface to colonize.

We showed that both WT and Δ*parDE_4_* strains react in a similar manner to the presence of excess eDNA (Fig. S2), so how can we reconcile this with our observation that WT cells disperse more efficiently than Δ*parDE_4_* cells (Fig. 4C)? *C. crescentus* biofilms are composed of discrete, clonal microcolonies (Fig. 4A and (Entcheva-Dimitrov & Spormann, 2004, Rossy *et al*., 2019). These microcolonies provide a microenvironment where conditions can differ from other microcolonies in the same biofilm. As cells die in biofilm microcolonies, the released eDNA will be at maximum concentration in the microcolony where it was released and will interact with nearby cells before diffusing to adjacent microcolonies where its concentration will be lower. Since eDNA inhibition of swarmer cell adhesion is proportional to its concentration, microcolonies with increased death rate will contain more eDNA and have a higher swarmer cell dispersal rate. Cells from WT microcolonies, which have a higher death rate in mature biofilms, should therefore have a higher rate of dispersal than cells from Δ*parDE_4_* microcolonies.

In addition to promoting dispersal, the PCD mechanism we describe here may also help preserve the initial biofilm. Indeed, once irreversibly attached via their holdfasts, stalked cells cannot disperse (Berne *et al*., 2010). In addition to stimulating dispersal of swarmer cells, PCD of some of the stalked cells in the biofilm may help reduce overcrowding, O_2_ starvation, and the death of the entire colony. Moreover, O_2_ limitation can result not only from overcrowding but also from environmental changes. Since *C. crescentus* is an obligate aerobe, this PCD mechanism can also act as a safety trigger that promotes the propagation of daughter cells to more hospitable environments when the colony starts to experience suboptimal O_2_ levels resulting from an external modification of the environment. Since the magnitude of inhibition of swarmer cell adhesion is proportional to eDNA concentration, this PCD mechanism may act as a rheostat tying cell dispersal to the availability of O_2_. Other PCD mechanisms in *C. crescentus* could similarly tie cell dispersal from the biofilm to other environmental conditions.

The role of PCD in biofilm regulation is well documented in different bacterial species and different TAS have been reported to play a role in biofilm regulation (Wang & Wood, 2011, Wen *et al*., 2014, Kamruzzaman *et al*., 2021, Singh *et al*., 2021). In *Escherichia coli*, several TAS are involved in regulation of biofilm formation at different stages (Kim *et al*., 2009, Kolodkin-Gal *et al*., 2009, Yamaguchi & Inouye, 2011, Soo & Wood, 2013, Zhao *et al*., 2013). Other TAS have been reported to regulate biofilms in different species, such as in *Vibrio cholerae* (Wang *et al*., 2015), *Staphylococcus aureus* (Kato *et al*., 2017), or *Pseudomonas aeruginosa* (Wood & Wood, 2016, Shmidov *et al*., 2022).

O_2_ limitation is one of the major environmental stresses experienced by bacteria in a mature biofilm (Stewart & Franklin, 2008, Kostakioti *et al*., 2013) and some species, such as *Shewanella oneidensis* or *P. aeruginosa*, respond to O_2_ depletion by triggering single cell dispersion (Thormann *et al*., 2005, Barraud *et al*., 2009, Rumbaugh & Sauer, 2020). PCD is involved in biofilm regulation upon O_2_ limitation in *S. aureus*, via the coordinated increased expression of the AltA murein hydrolase and decreased expression of the cell wall-teichoic acids (Mashruwala *et al*., 2017). In addition, the Hha/TomB TAS regulates cell death in *E. coli* biofilms in response to O_2_ availability (Marimon *et al*., 2016): under high O_2_ conditions, the toxin is inactivated by spontaneous oxidation enhanced by transient interactions with the antitoxin under those conditions. In O_2_ limited regions of the biofilm, oxidation of the Hha toxin stimulates cell death and enables the formation of gaps in the biofilm matrix, thereby facilitating biofilm dispersal through those gaps (García-Contreras *et al*., 2008, Marimon *et al*., 2016). However, to our knowledge, none of the previously reported systems function as described in this study.

We showed that the O_2_-limited conditions where *parDE_4_*-dependent cell death occurs correspond to the lowest level of *parDE_4_* transcription. This result suggests the possibility that lowering the expression of the *parDE_4_* operon triggers cell death. During surface adhesion and in early stages of biofilm development, O_2_ availability is good relative to later stages when biomass increases, O_2_ becomes limiting, and cell death is increased. It is interesting to note that the “neutralized” steady state of the ParDE_4_ TAS, when the toxin is inactivated, seems to be when the operon transcription is at its highest, i.e, when O_2_ is abundant. Indeed in most studied TAS, neutralization of the toxin is linked to a decrease in transcription of the operon, and stresses have been shown to induce transcription of TAS (LeRoux *et al*., 2020, Jurėnas *et al*., 2022). We are unaware of cases where reduced TAS expression is correlated with the condition that activates the PCD in biofilm regulation. This suggests a novel regulatory mechanism of PCD that remains to be understood.

TAS have often been reported to play a direct and crucial role in adaptation to different stresses. However, the physiological role of TAS-regulated cell death has been challenged in recent years (Harms *et al*., 2017, Goormaghtigh *et al*., 2018, Fraikin *et al*., 2019, Wade & Laub, 2019, Fraikin *et al*., 2020, Rosendahl *et al*., 2020). Even if a TAS operon is transcribed under stress conditions, it does not necessarily mean that the toxin is liberated, and transcription might not always reflect TAS activity (LeRoux *et al*., 2020). Therefore, whether the decrease in *parDE_4_* transcription that correlates with increased cell death is mechanistically related to TA activation is still not known.

Our future work will focus on identifying the pathway that controls the activation of the ParDE_4_ mediated PCD. Based on sequence similarity, ParDE_4_ belongs to the type II TAS family (Fiebig *et al*., 2014). In *C. crescentus*, the ParDE_4_ homolog ParDE_1_ is thought to have a bacteriostatic effect, inhibiting cell division and causing cell filamentation by an unknown mechanism (Fiebig *et al*., 2014). In *E. coli*, the plasmid borne ParDE TAS inhibits DNA replication by targeting DNA gyrase, leading to cell death (Jiang *et al*., 2002). However, *E. coli* ParE_2_ toxin, whose structure is close to that of the predicted *C. crescentus* ParE_1_ (Dalton & Crosson, 2010) does not bind to DNA gyrase and the molecular target of this toxin is still unknown (Sterckx *et al*., 2016). These results suggest that the ParDE family is not homogenous in terms of targets, mechanisms, and physiological outcomes.

Does the PCD-stimulated cell dispersal mechanism we have described constitute cooperation and even altruism? Altruism mechanisms in biofilms have been discussed previously (Penesyan *et al*., 2021, Sadiq *et al*., 2021). Altruism involves a trade-off between growth rate and growth yield in the biofilm to preserve resources at the cost of individual growth rate for the fitness of the entire population (Kreft, 2004), or with the public goods that are used by the whole attached colony (Nadell *et al*., 2008). In addition, to be beneficial for the overall community while being costly for the individual, Hamilton’s rule states that altruism must promote only the success of the surviving kin (Hamilton, 1970, Ramisetty & Sudhakari, 2020). We showed previously that eDNA stimulation of dispersal through holdfast inhibition is highly specific to *Caulobacter* species: biofilms of other tested species were not impaired by addition of eDNA, including *Caulobacter* DNA, and *Caulobacter* adhesion is only inhibited by eDNA from *Caulobacter* (Berne *et al*., 2010). In addition, our current results show that deletion of ParDE_4_ decreases dispersal from the mature biofilm. Intuitively, stimulating dispersal away from a biofilm where environmental quality is declining to another, potentially better environment would appear to be advantageous for the dispersing kin in terms of fitness, but this has not been directly demonstrated in our case. In addition, reducing the competing population in a mature biofilm with declining O_2_ availability could provide an advantage to the surviving attached cells, but this is difficult to determine experimentally. Thus, the PCD-mediated mechanism described in this study might confer a fitness advantage to the overall *Caulobacter* population under the appropriate environmental conditions.

## MATERIALS AND METHODS

### Bacterial strains and growth conditions

All bacterial strains used in this study are listed in Table S1. *Escherichia coli* strains used for cloning experiments were grown in LB medium at 37°C. When necessary, antibiotics were added at the following final concentrations: kanamycin (Kan) 50 µg ml^-1^ (for pMT630 constructs), and tetracycline (Tet) 15 µg ml^-1^ (for pRKlac290 constructs). *C. crescentus* strains were grown at 30°C using M2 minimal medium complemented with 0.2% glucose (M2G) (Johnson & Ely, 1977) in liquid, and using Peptone Yeast Extract (PYE) (Poindexter, 1964) + 15 g l^-1^ bactoagar (Difco) plates. *C. crescentus* cultures were grown under four different aeration conditions: (1) maximal aeration by growing 1 ml of culture in a 15 ml glass culture tube tilted and shaken at 300 rpm; (2) high aeration with 1 ml of culture in a 2 ml microtube sealed with a 1.5 x 1.5 cm piece of sterile breathable sealing film (AeraSeal, Excel Scientific), to allow air-exchange, in a shaking ThermoMixer (Eppendorf), at 100 rpm; (3) moderate aeration with 1 ml of a culture in a 2 ml microtube sealed with a 1.5 x 1.5 cm piece of sterile breathable sealing film (AeraSeal, Excel Scientific), to allow air-exchange, in a non-shaking ThermoMixer (Eppendorf); and (4) limited aeration with 1 ml of a culture in a 2 ml microtube tightly sealed with an air tight rubber plug to prevent air exchange with the environment, in a non-shaking ThermoMixer (Eppendorf). When appropriate, Kan and Tet were added to 20 and 2 µg ml^-1^ when using plasmids pMT630 and pRKlac290, respectively. To grow the pMR10-P*ccoN*-*mcherry* pMR20-P*parDE_4_*-*gfp* harboring strain, both Kan and Tet were added to 0.5 µg ml^-1^.

*C. crescentus ΔparDE_4_* strains harboring the stable miniTn7*gfp* or miniTn7*dsred* (CB15 Δ*parDE_4_*::miniTn7*gfp* (YB5253) and CB15 Δ*parDE_4_*::miniTn7*dsred* (YB5254)) were constructed by ϕCr30 mediated transduction (West *et al*., 2002) from AS110 and AS109 (Entcheva-Dimitrov & Spormann, 2004), respectively, into *C. crescentus* CB15 Δ*parDE_4_* (FC915).

### Plasmid construction and cloning procedures

All plasmids were cloned using standard molecular biology techniques. PCR were performed using *C. crescentus* CB15 WT (YB135) gDNA as the template. Sequences of the primers used are available upon request.

To construct a *parD_4_* inducible expression plasmid, the entire *parD_4_* gene was cloned in frame into the low copy vanillate-inducible replicating plasmid pMT630 (Thanbichler *et al*., 2007), to give plasmid pMT630-*parD_4_*. To construct a transcriptional fusion plasmid for the *parDE_4_* promoter, 567 bp upstream of the *parD_4_* start codon were cloned into pRKlac290 (Gober & Shapiro, 1992) upstream and in the frame of *lacZ*, to give plasmid pRKlac290-P*parDE_4_*.

### Epifluorescence microscopy

Epifluorescence microscopy was performed using an inverted Nikon Ti2 microscope with a Plan Apo 60X objective, a GFP/DsRed filter cube, an Andor iXon3 DU885 EM CCD camera, and Nikon NIS Elements imaging software. Image analysis was performed using ImageJ functions and plug-ins (Schneider *et al*., 2012).

### Live/dead quantification in planktonic cultures

Cell death in planktonic cultures was detected using the BacLight Bacterial Viability kit (L7007, Invitrogen) and quantified by fluorimetry. A ratio of 1:1 Live/Dead stain mixture from the kit was diluted to 1/1000 in sterile dH_2_O. One hundred µl of diluted Live/Dead stain were added to 100 µl of the liquid culture to be assayed. The resulting 200 µl were pipetted to a black 96-well plate with a clear flat bottom (Corning) prior to measurements using a Synergy HT microplate reader. Standard samples were added to each plate of samples, to generate a calibration curve that allows experimental samples quantification. Mid-log phase *C. crescentus* cells were diluted to an OD_600_ of 0.1. The diluted culture was divided into two samples: one sample was mixed with an equal volume of M2 medium (with no carbon source) (Johnson & Ely, 1977), to become the live cell standard sample; and one sample was mixed with an equal volume of 70% isopropyl alcohol, to become the dead cell standard sample. Different ratios of live and dead cell samples were mixed to provide five standards (ratio live:dead of 100:0, 75:25, 50:50, 25:75, 0:100). Absorbance at 600 nm, green and red fluorescence signals (using 485/528 nm and 485/630 nm Em/Ex filters, respectively) were quantified for each well. Data were analyzed by calculating the ratio of Fluorescence emission at 528 nm (green signal) / Fluorescence emission at 630 nm (red signal). The values calculated for the standard samples were plotted versus the percentage of live cells in those standards, and a linear regression from the standard curve was used to determine the percentages of live and dead cells in each sample.

### eDNA quantification

eDNA was quantified using the Quant-iT PicoGreen reagent (Invitrogen), as described previously (Berne *et al*., 2010). Briefly, PicoGreen was diluted 1:200 in TE (10 mM Tris-HCl, 1 mM EDTA, pH 7.5) buffer prior analysis. A volume of 150 µl of planktonic phase was centrifuged (2 min at 15,000 g, RT) to pellet cells and 100 µl of supernatant was added to 100 µl diluted PicoGreen reagent in a black 96-well plate with a clear flat bottom (Corning). After 5 minutes of incubation at room temperature, the fluorescence was measured using a Synergy HT microplate reader (using 485/528 Ex/Em nm filter set). A calibration curve (using Lambda DNA standard provided in the Quant-it kit, from 0 to 20 µg ml^-1^) was performed for each measured plate. The fluorescence values measured for the standard samples were plotted versus the DNA concentration in those standards, and the linear regression from the standard curve was used to determine the DNA concentration from each sample.

### Biofilm quantification by crystal violet staining

*C. crescentus* cells were grown to mid-log phase in M2G medium under shaking conditions and diluted to OD_600_ = 0.1 in the same culture medium. Aliquots of 1 ml of diluted cells were placed in the aforementioned growth conditions consisting of various aeration levels. After incubation at 30°C, the planktonic phase was carefully removed from the tube. The tube was rinsed with sterile dH_2_O to remove non-attached cells, and the biomass attached to the inside of the tube was stained with a 0.1% crystal violet solution for 5 min and rinsed again with sterile dH_2_O to remove excess crystal violet. The crystal violet stained cells were eluted with 10% acetic acid and quantified by measuring A_600_.

### Spent medium preparation

Spent medium was prepared as previously described (Berne *et al*., 2010). Briefly, bacteria were grown for 36 h (late stationary phase) at 30°C in M2G medium. Cells were removed from the cultures by centrifugation (10 min at 8,000 g, RT) and the supernatant (spent medium) was filter-sterilized using a 0.2 µm filter and kept at −20°C until used.

### Planktonic cells visualization by microscopy

Exponential phase cultures (OD_600_ = 0.4-0.6) were spotted on a glass coverslip (1 µl) and covered by a 1% agarose pad in M2G (Seakem LE agarose). Epifluorescence microscopy was performed as described below.

### Biofilm visualization by microscopy

For microscopy analysis, biofilms were grown in 2 ml microtubes under moderate aeration conditions as described above, with the only difference that a 22 x 5 mm strip of PVC coverslips (ref 12-547, Fisher Scientific) placed vertically in the microtube before sealing with Aeraseal. After incubation at 30°C, the coverslip was rinsed with sterile dH_2_O to remove non-attached cells and stained for microscopy as described below (Live/dead staining or FDG labeling).

### Live/Dead staining of cells inside the biofilm and quantification using microscopy images

Rinsed biofilm covered PVC strips (see above) were stained with the BacLight Bacterial Viability kit (L7007, Invitrogen). A volume of 0.5 µl of 1:1 Live/Dead stain mixture was added to 500 µl sterile dH_2_O, placed on top of the strip and incubated for 15 min at room temperature in the dark. The stained biofilm was then rinsed and observed by epifluorescence microscopy. Live cells quantification was performed on microscopy images obtained above as described previously (Berne *et al*., 2010). Fluorescent signals coming from the green (live cells) and the red (dead cells) channels were quantified using the ImageJ analysis software (Schneider *et al*., 2012). 16-bit images were thresholded using the B/W default setting and the total area of fluorescent signal was automatically determined using the Analyze Particle function, as the fraction area. For a given picture, the areas for the green and red signals were added and adjusted to 100% and percentage of each signal was then calculated. Results are given as the average from 10 random images for each biofilm sample, performed in independent triplicates.

### FDG staining

The activity of the *parDE_4_* promoter in the biofilm was monitored using the plac290-P*parDE_4_* construct and the fluorescein Di-β-D-Galactopyranoside (FDG), a fluorescent substrate for β-galactosidase (Rotman *et al*., 1963). Rinsed PVC strips (see above) covered with biofilms were stained with FDG (Invitrogen) using the following procedure. Firstly, strips were rinsed with sterile dH_2_O. Secondly, 1 µl of FDG was added to 100 µl sterile dH_2_O, placed on top of the strip and incubated for 15 min at room temperature in the dark. Finally, the stained biofilm was rinsed again and observed by epifluorescence microscopy. Fluorescence quantification was performed using the ImageJ analysis software (Schneider *et al*., 2012). Intensity is defined as the average signal value measured on the detected particle. Results are given as the average intensity from 10 random images for each biofilm sample, performed in independent triplicates.

### Competition experiments in flow-cells

Biofilm were grown in flow-cells using the previously described set-up (Berne *et al*., 2010). Mid-log phase cultures of WT and Δ*parDE*_4_ containing the mini*Tn7*::*gfp* or mini*Tn7*::*dsred* insertion were diluted to an OD_600_ of 0.025 and mixed to 1:1 ratio prior inoculation in the flow cell (200 µl). Initial attachment was performed in the absence of flow for 1 h, followed by a constant flow of 3 ml h^-1^. Surface colonization of the glass surface covering the flow cells was monitored for seven days at room temperature. Epifluorescence images were recorded at different time points, the amount of each population was quantified (as the fluorescence area for green and red signals), and the ratio of each population was calculated (as a percentage of the total population for the given time point). Each experimental condition was performed twice (swapping GFP and dsRed labelling for each replicate) in parallel triplicate chambers.

### Cell dispersal quantification

The amount of cells released from the biofilm grown in a flow-cell (as described above) was determined by counting the number of colony forming units (CFU) in the flow through. A sample of 1 ml was collected 5 cm downstream of the flow cells. CFU were determined by generating 1:5 serial dilution of the flow through in M2G. Twenty µl of each dilution were spotted on a dry PYE-agar plate, in triplicate, and incubated at 30°C for 2 days. Only spots of serial dilutions which contained 5 to 30 colonies were considered for analysis. Plates were imaged using a ChemiDoc imaging system (Biorad), using the 530 nm and 605 nm emission fluorescence filter reading parameters, to allow quantification of GFP and DsRed labelled colonies in each spot.

### ß-galactosidase promoter activity assays

ß-galactosidase activity was quantified colorimetrically in 96-well plates as described previously (Miller, 1972, Berne *et al*., 2018, Berne & Brun, 2019). Overnight cultures of strains bearing plac290lacZ or its derivatives were diluted to OD_600_ of 0.05 and incubated at 30°C until an OD_600_ of 0.4 – 0.6 was reached. A culture volume of 200 µl was mixed with 600 µl of Z buffer (60 mM Na_2_HPO_4_, 40 mM NaH_2_PO_4_, 10 mM KCl, 1 mM MgSO_4_, 50 mM ß-mercaptoethanol). Cells were then permeabilized using 50 µl of chloroform and 25 µl of 0.1% SDS. Two hundred µl of the substrate *o*-nitrophenyl-β-D-galactoside (4 mg ml^-1^) were added to the permeabilized cells. Upon development of a yellow color, the reaction was stopped by raising the pH to 11 with addition of 400 µl of 1M Na_2_CO_3_. Absorbance at 420 nm (A_420_) was determined and the Miller Units of β-galactosidase activity were calculated as (A_420_)(1000)/(OD_600_)(*t*)(*v*) where *t* is the time in minutes and *v* is the volume of culture used in the assay in ml. The β-galactosidase activity of CB15 plac290 (empty vector control) was used as a blank sample reference.

### Quantitative PCR

WT and Δ*parDE_4_* strains were grown overnight under the four aeration conditions described above, then diluted to OD_600_ of 0.05 and incubated at 30°C under the same aeration until an OD_600_ of 0.4 – 0.6 was reached. Cells were centrifuged at 4°C and pellets were stored at −80°C until further use. RNA samples were prepared using the Monarch Total RNA Miniprep kit (New England Biolabs Inc.), followed by a DNAse treatment using the turbo DNA-free kit (Life Technologies), and storage at −80°C. One-step quantitative RT-PCR was used to determine gene expression levels using the Luna Universal One-Step RT-qPCR kit (New England Biolabs Inc.) according to the manufacturer’s recommendations using 4 ng of total RNA per 20µl reaction. The reactions were performed on a QuantStudio 3 device (Applied Biosystems, Thermo Fisher Scientific). Gene expression ratios were calculated using *rpoD* (CCNA_03142) as the reference gene and the *ccoN* (CCNA_01467) as the target gene. Sequences of the primers used were as follow: rpoD-for (GCAGCTCTATGCGATCAACA), rpoD-rev (TGTCGTTCTCGACGAACTTG), ccoN-for (TGGCCGACGATCTTCTACTT), ccoN-rev (AACAGCTGATAGCCCCAGAA). Each gene expression value was obtained from 2 independent biological replicates, grown on separate days. RT-qPCR reactions were run in triplicate. Gene expression ratios were calculated using the Pfaffl method (Pfaffl, 2001), which takes into account primer efficiencies.

## ACKNOWLEDGEMENTS

The authors thank Aretha Fiebig and the Crosson laboratory for providing strains and engaging discussions, as well as Marylise Duperthuy for the use of the QuantStudio 3 apparatus, and the members of the Brun laboratory for their constant feedback. This study was supported by grant R35GM122556 from the National Institutes of Health and by a Canada 150 Research Chair in Bacterial Cell Biology to YVB.

---

**Figure S1: ParDE4 is involved in eDNA release under moderate aeration conditions.** (A) Representative growth curves of cultures grown at 30°C, followed by OD_600_ over time. Cultures were grown under moderate aeration (no shaking, 1 ml culture in a 2 ml microtube sealed with AeraSeal breathable film), and under maximal aeration conditions (300 rpm shaking, 1 ml culture in a 15 ml culture tube). (B) Quantification of eDNA released in the planktonic phase in the same cultures. Results are given as the average of at least three independent experiments and the error bars represent the Standard Error of the Mean (SEM).

**Figure S2: Biofilm inhibition in WT and Δ*parDE_4_* by eDNA from different spent media.** Biofilm formation of WT (gray circles) and Δ*parDE_4_* (white squares) at different total eDNA concentrations. Because Δ*parDE_4_* produces less eDNA than WT, we first determined the amount of eDNA present in each culture and added an appropriate amount of spend medium to obtain the same final concentration of eDNA. Results are given as a percentage of WT without exogenous addition of eDNA, of three independent experiments, each run in duplicate.

**Figure S3: Expressions of the *parDE_4_* operon is reduced when a hypoxia reporter expression is increased.** Red and green fluorescence intensity of single cells grown under maximal (A), high (B), moderate (C), and limited (D) aeration. WT cells carrying both pMR20-P*parDE_4_-gfp* and pMR10-P*ccoN-mcherry* plasmids were grown to an OD_600_ of 0.4 to 0.6 and imaged by epifluorescence microscopy. Fluorescence intensity was calculated from more than 5,000 cells from at least three independent replicates.

**Table S1:** Plasmids and strains.

|  | Description / Construction | Source / Reference |
| --- | --- | --- |
| **Strains** |  |  |
| YB135 | CB15 WT | (Poindexter, 1964) |
| YB4789 | CB15 WT::miniTn7-*gfp* | (Berne *et al.*, 2010) |
| YB4788 | CB15 WT::miniTn7-*dsred* | (Berne *et al.*, 2010) |
| AS110 | CB15 WT::miniTn7-*gfp* | (Entcheva-Dimitrov & Spormann, 2004) |
| AS109 | CB15 WT::miniTn7-*dsred* | (Entcheva-Dimitrov & Spormann, 2004) |
| YB5253 | CB15 Δ*parDE*<sub>4</sub>::miniTn7-*dsred* | This study |
| YB5254 | CB15 Δ*parDE*<sub>4</sub>::miniTn7-*dsred* | This study |
| FC304 | CB15 Δ*parDE*<sub>1</sub> | (Fiebig *et al.*, 2010) |
| FC975 | CB15 $\Delta$ Ψ*parDE*<sub>2</sub> | (Fiebig *et al.*, 2010) |
| FC922 | CB15 Δ*parDE*<sub>3</sub> | (Fiebig *et al.*, 2010) |
| FC915 | CB15 Δ*parDE*<sub>4</sub> | (Fiebig *et al.*, 2010) |
| FC917 | CB15 Δ*relBE*<sub>1</sub> | (Fiebig *et al.*, 2010) |
| FC993 | CB15 Δ*relBE*<sub>2</sub> | (Fiebig *et al.*, 2010) |
| FC980 | CB15 Δ*relBE*<sub>3</sub> | (Fiebig *et al.*, 2010) |
| FC977 | CB15 Δ*relBE*<sub>4</sub> | (Fiebig *et al.*, 2010) |
| | CB15 Δ*parDE* | Aretha Fiebig |
| | CB15 Δ*relBE* | Aretha Fiebig |
| | CB15 Δ*parDE relBE* | Aretha Fiebig |
| FC878 | CB15 Δ*parE*<sub>4</sub> | (Fiebig *et al.*, 2010) |
| YB8243 | CB15 WT pRKlac290 | (Berne *et al.*, 2018) |
| YB7040 | CB15 WT pRKlac290-*PparDE*<sub>4</sub> | This study |
| YB9699 | CB15 WT pRKlac290-*PccoN* | This study |
| YB8944 | CB15 WT pMT630 | (Berne *et al.*, 2018) |
| YB9408 | CB15 WT pMT630-*parD*<sub>4</sub> | This study |
| YB9409 | CB15 Δ*parE*<sub>4</sub> pMT630 | This study |
| YB9410 | CB15 Δ*parE*<sub>4</sub> pMT630-*parD*<sub>4</sub> | This study |
| YB5282 | CB15 WT pMR10-*PccoN-mcherry* | This study |
| YB4779 | CB15 WT pMR20-*PparDE*<sub>4</sub>-*gfp* | This study |
| YB9491 | CB15 WT pMR20-*PparDE*<sub>4</sub>-*gfp* pMR10-*PccoN-mcherry* | This study |
| **Plasmids** |  |  |
| pRKlac290 | plasmid for *lacZ* transcriptional fusions | (Gober & Shapiro, 1992) |
| pRKlac290-*PparDE*<sub>4</sub> | *parDE*<sub>4</sub> promoter region cloned into pRKlac290 | This study |
| pRKlac290-*PccoN* | *ccoN* promoter region cloned into pRKlac290 | This study |
| pMT630 | Low copy replicating plasmid, vanillate inducible, Kn resistant | (Thanbichler *et al.*, 2007) |
| pMT630-*parD*<sub>4</sub> | *parD*<sub>4</sub> under control of the vanillate promoter in pMT630 | This study |
| pMR20 | Replicating plasmid, IPTG inducible (constitutive in *C. crescentus*) (Tet resistant) | R. Roberts |
| pMR20-*PparDE*<sub>4</sub>-*gfp* | *gfp* expression under the control of the *parDE*<sub>4</sub> promoter | This study |
| pMR10 | Replicating plasmid, IPTG inducible (constitutive in *C. crescentus*) (Kn resistant) | R. Roberts |
| pMR10-*PccoN-mcherry* | *mCherry* expression under the control of the *ccoN* promoter | This study |

