## Supplementary figures and images for "eDNA-stimulated cell dispersion from *Caulobacter crescentus* biofilms upon oxygen limitation is dependent on a toxin-antitoxin system"

### Supplementary figures

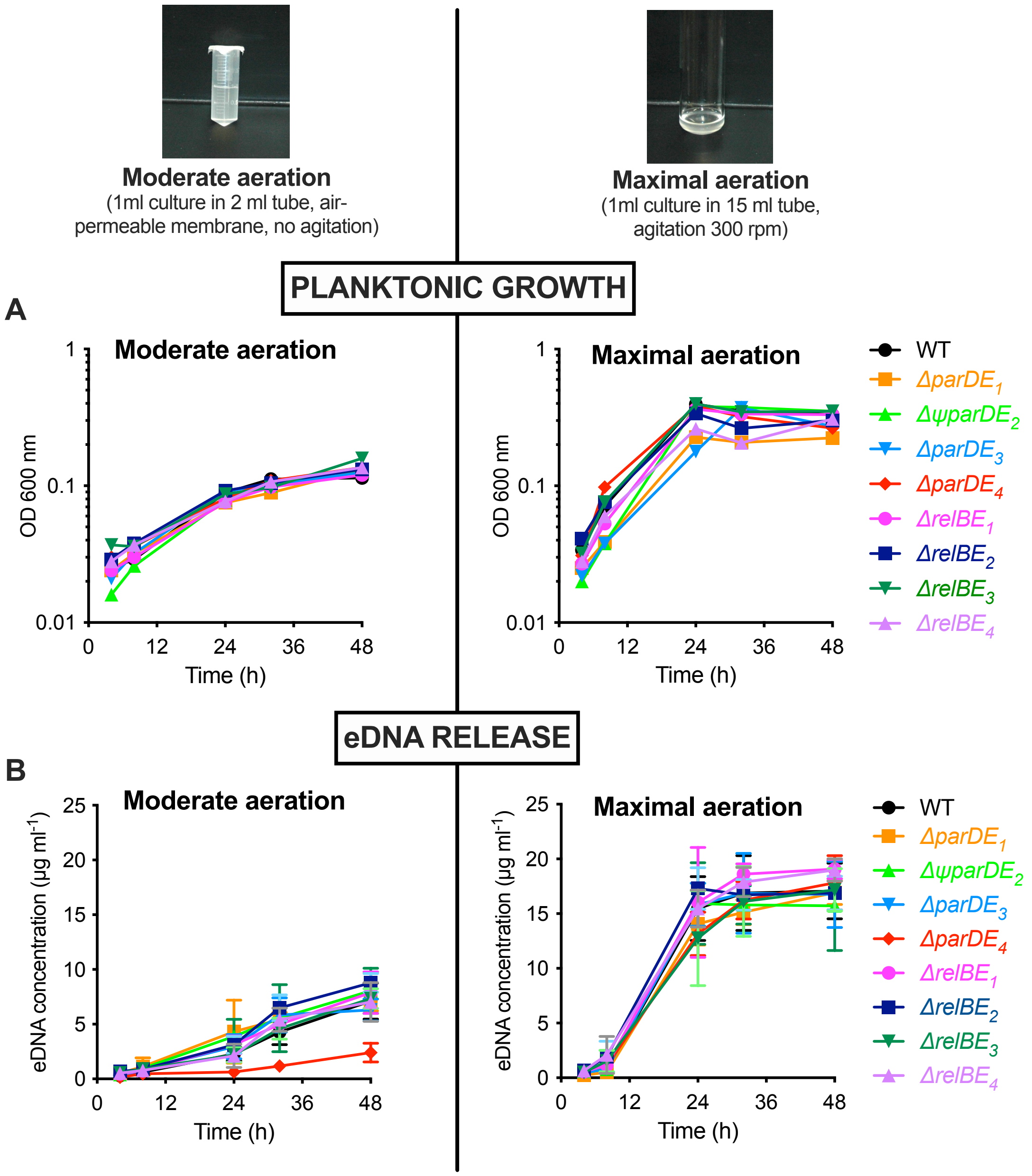

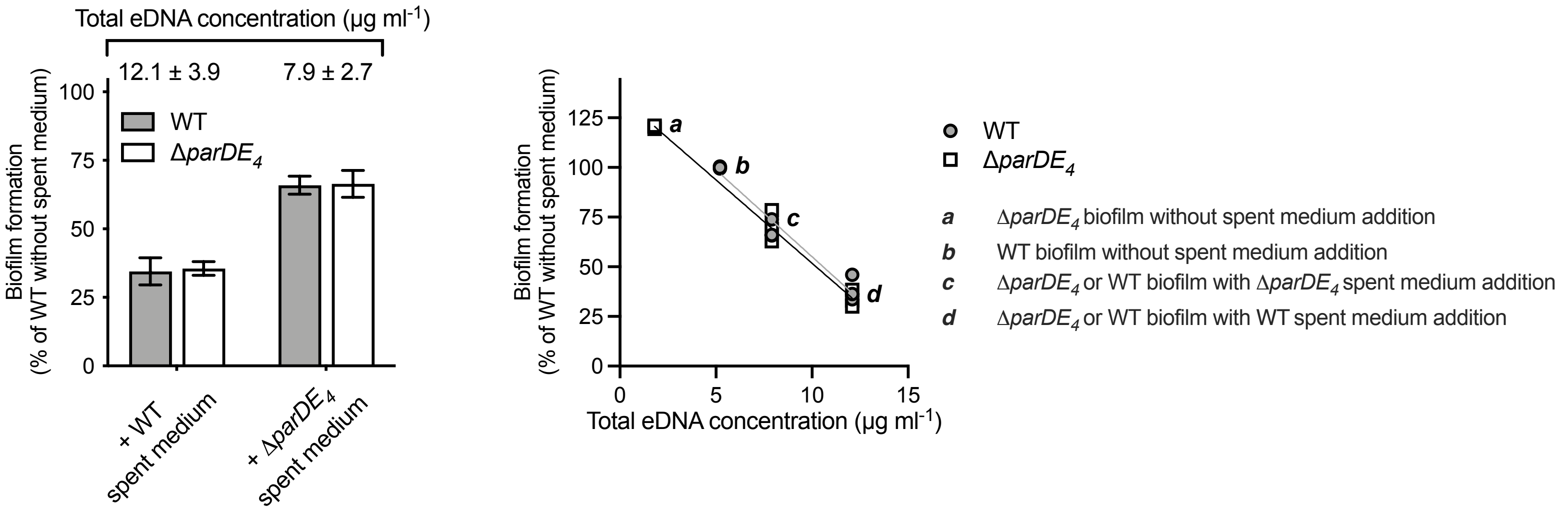

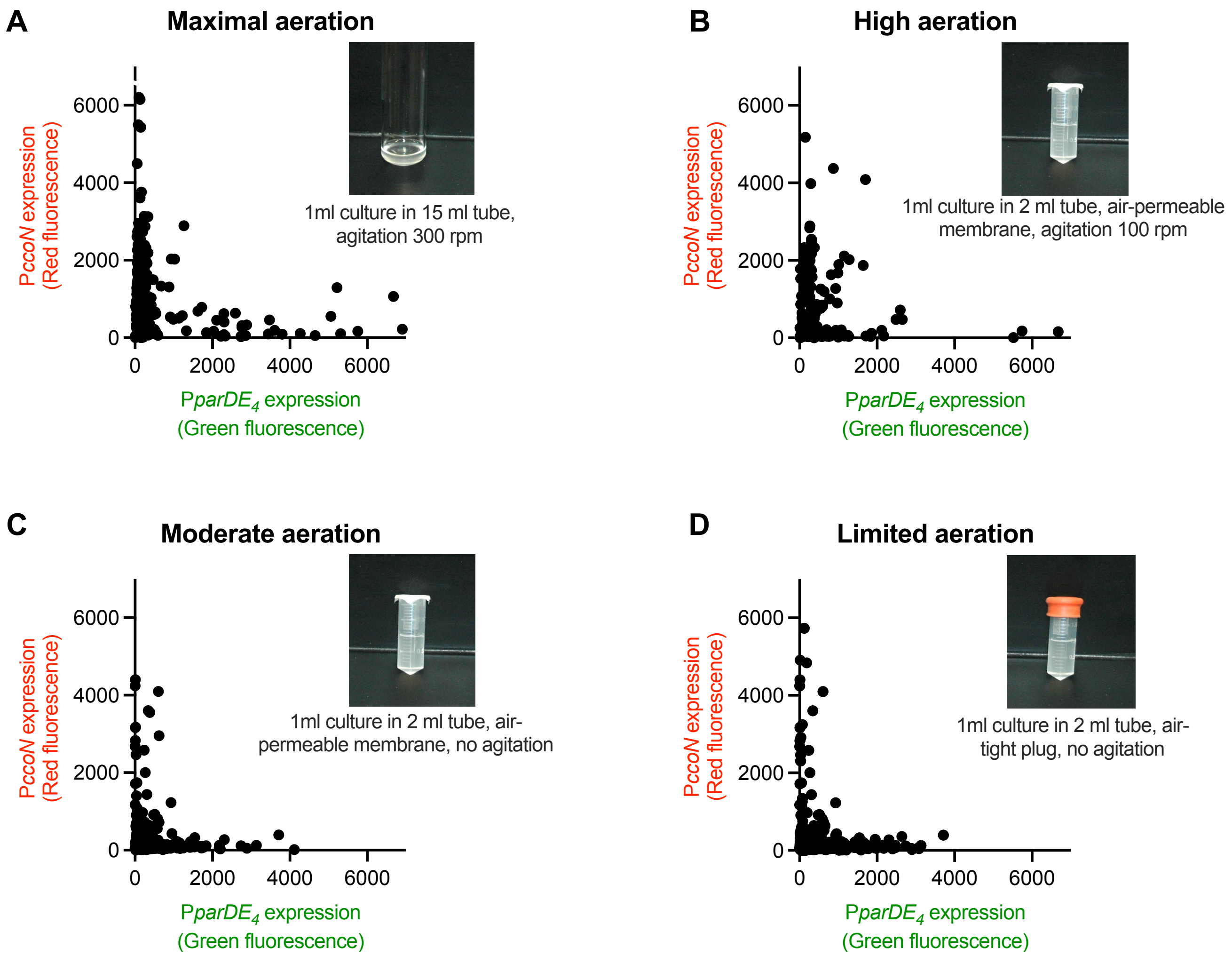
